# Protease Activity Analysis: A Toolkit for Analyzing Enzyme Activity Data

**DOI:** 10.1101/2022.03.07.483375

**Authors:** Ava P. Soleimany, Carmen Martin Alonso, Melodi Anahtar, Cathy S. Wang, Sangeeta N. Bhatia

## Abstract

Analyzing the activity of proteases and their substrates is critical to defining the biological functions of these enzymes and to designing new diagnostics and therapeutics that target protease dysregulation in disease. While a wide range of databases and algorithms have been created to better predict protease cleavage sites, there is a dearth of computational tools to automate analysis of *in vitro* and *in vivo* protease assays. This necessitates individual researchers to develop their own analytical pipelines, resulting in a lack of standardization across the field. To facilitate protease research, here we present Protease Activity Analysis (PAA), a toolkit for the preprocessing, visualization, machine learning analysis, and querying of protease activity datasets. PAA leverages a Python-based object-oriented implementation that provides a modular framework for streamlined analysis across three major components. First, PAA provides a facile framework to query datasets of synthetic peptide substrates and their cleavage susceptibilities across a diverse set of proteases. To complement the database functionality, PAA also includes tools for the automated analysis and visualization of user-input enzyme-substrate activity measurements generated through *in vitro* screens against synthetic peptide substrates. Finally, PAA can supports a set of modular machine learning functions to analyze *in vivo* protease activity signatures that are generated by activity-based sensors. Overall, PAA offers the protease community a breadth of computational tools to streamline research, taking a step towards standardizing data analysis across the field and in chemical biology and biochemistry at large.

## Introduction

Proteases play essential roles in diverse biological processes ranging from development to differentiation, and dysregulated protease activity is a driver of a variety of pathological conditions including cancer, neurodegeneration, and infectious diseases.^1^ Because proteases most proximally exert their function through their *activity*, understanding protease activity, rather than transcript or protein expression, is required to elucidate the biological roles of proteases and to harness these enzymes as diagnostic and therapeutic targets. To this end, molecular tools such as activity-based probes (ABPs), short synthetic peptide substrates, and noninvasive enzyme activity sensors have been developed to measure protease activities *in vitro*, i.e., of recombinant proteases or enzymes present in biospecimens, ^2–6^ as well as *in vivo*, i.e., within the disease microenvironment.^7–11^ Beyond their use as a discovery tool, sensors that quantify protease activity are being applied directly for early detection and monitoring of disease,^5,12–17^ biological imaging, ^7,18^ and drug discovery.^19,20^ Furthermore, proteolytic cleavage of peptide linkers is being used to trigger disease-specific activation of engineered activity-based diagnostics^11^ and therapeutics,^21–24^ all of which inherently rely on assessments of protease activity for their design and optimization. To support the development of these new activity-based tools and to advance the study of protease biology at large, a wide range of databases and algorithms have been created to better identify protease substrates and cleavage sites, providing a clear demonstration of how protease research can benefit from computational tools.^25–27^

Rapidly identifying, designing, and characterizing new peptide substrates and activity-based sensors remains a major bottleneck towards advancing these applications, due to the promiscuous nature of protease activity, the combinatorial number of substrates synthetically accessible, and the dearth of methods to automate protease activity analysis and substrate design. Current protease databases also focus exclusively on endogenous substrates and cleavage sites,^28^ despite the fact that synthetic activity-based sensors and large-scale libraries of synthetic peptides are now standard tools for measuring and quantifying protease activity *in vivo* and *in vitro*. Furthermore, the development of these experimental and molecular methods has not been accompanied by scalable, modular data analytic workflows. As a result, individual researchers and their affiliated labs are constantly re-inventing the wheel when it comes to analyzing their data. The creation of computational packages similar to what exists for genomics (e.g., Bioconductor) could enable the standardized analysis of data generated from both *in vitro* and *in vivo* protease-based experiments. This would greatly benefit researchers by accelerating experimental workflows, informing diagnostic and therapeutic design, and facilitating biological insight into protease dysregulation in disease.

To address these needs, we present Protease Activity Analysis (PAA), a toolkit that addresses the need for computational methods to accelerate data analysis of enzyme activity data in biochemistry, chemical biology, and bioengineering (Fig. 1). PAA contains a searchable database of existing protease activity data, curated from over a decade of published works from our group, along with modular analytics that enable users to query these datasets for enzymes or substrates of interest. Through PAA’s framework, users can additionally create and search new databases using their own protease-substrate screening data. PAA enables analytic standardization via functions that automate the quantification and visualization of user-inputted data from *in vitro* protease activity screens and *in vivo* protease-activated nanosensors. The package is accompanied by step-by-step tutorials that detail the functionalities provided by PAA in an open source repository (https://github.com/apsoleimany/protease_activity_analysis). PAA’s Python-based implementation provides a modular framework that is easy to interface with other software packages and can be readily integrated into broader data analytic workflows.

**Figure 1:**
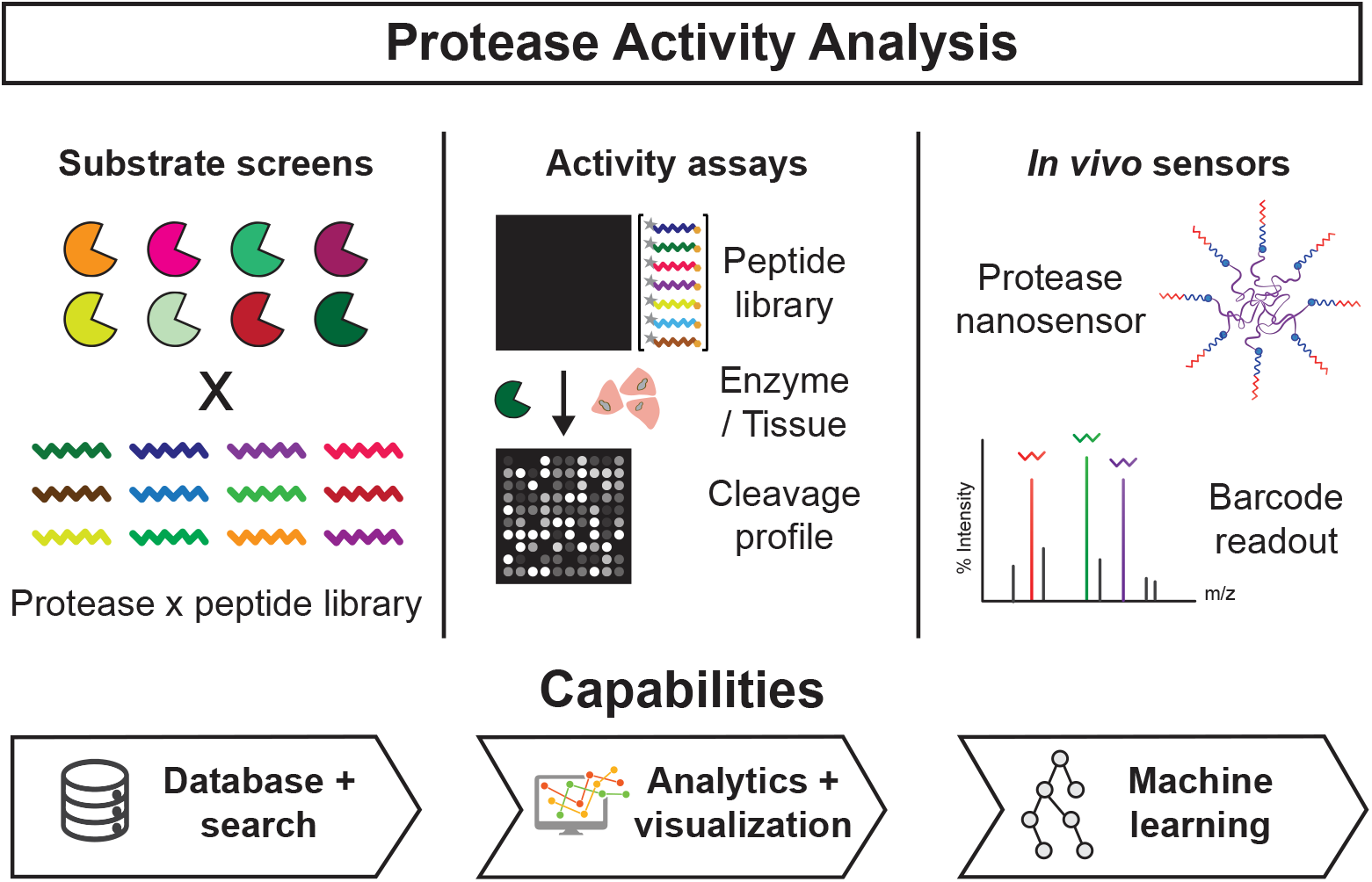
Overview of Protease Activity Analysis (PAA). The PAA package is designed to analyze data from large-scale substrate screens, enzyme activity assays, and *in vivo* enzyme activity sensors. Key package capabilities include searchable databases, where users can both query pre-loaded protease-substrate datasets published as part of PAA, or import new datasets privately for their own use; data analytic and visualization functions, for facile and automated analysis of protease activity data; and machine learning models, for classification analysis of activity-based sensor data.

## Results

`PAA` provides scalable and modular analysis capabilities for datasets of enzym activity measurements. Specifically, `PAA` supports three core analytic workflows (Fig. 1): (1) analysis and query of databases of peptide substrate sequences and their cleavage susceptibilities; (2) analysis of substrate screens using recombinant enzymes or biospecimens; (3) analysis of measurements from protease-responsive *in vivo* nanosensors. Across all three workflows, `PAA` provides preprocessing, visualization, machine learning, and search functionalities.

### PAA supports searchable enzyme-substrate databases

Identifying and characterizing which substrates are robustly and specifically cleaved by protease(s) of interest, for example those that are overexpressed in a specific cancer, is critical to discovery and engineering efforts that seek to understand and exploit protease activity. Indeed, the rapid rise and promise of engineered conditionally-activated diagnostics and therapeutics, which almost universally incorporate a protease-cleavable peptide linker as the “trigger” for disease-specific activation, has motivated the need for tools and methods that facilitate identification of synthetic peptide substrates maximally cleaved in diseased tissues and/or by target proteases. To this end, in `PAA` we present a `SubstrateDatabase` data structure that provides a facile framework for curating and querying datasets of enzyme-substrate activity, which often take the form of fluorometric assays of proteolytic cleavage of synthetic, fluorogenic peptide substrates. These assays measure the kinetics of enzyme activity over time and can be used to assess both the efficiency of an enzyme for a particular substrate, by quantifying the initial rate of the reaction, as well as the specificity of a substrate for a protease, by comparing the substrate’s cleavage against other proteases screened.

To demonstrate these capabilities, we have created a publicly available database that incorporates data generated by our group from 6 independent recombinant protease screens against fluorogenic peptide substrates. The database consists of 150 unique synthetic peptide substrates and their cleavage susceptibilities across a set of 77 distinct recombinant proteases spanning the metallo, serine, cysteine, and aspartic catalytic classes. The substrates published as part of PAA were identified based on the literature and designed to query the activity of disease-associated proteases, including those in cancer, infection, and thrombosis. As such, there exists an over-representation of metallo- and serine-sensitive substrates in the dataset (Fig. 2B).^13,29^ This dataset is open-sourced as part of PAA. In addition, users can import new data into PAA, instantiating `SubstrateDatabase` data structures that can be readily queried and analyzed.

**Figure 2:**
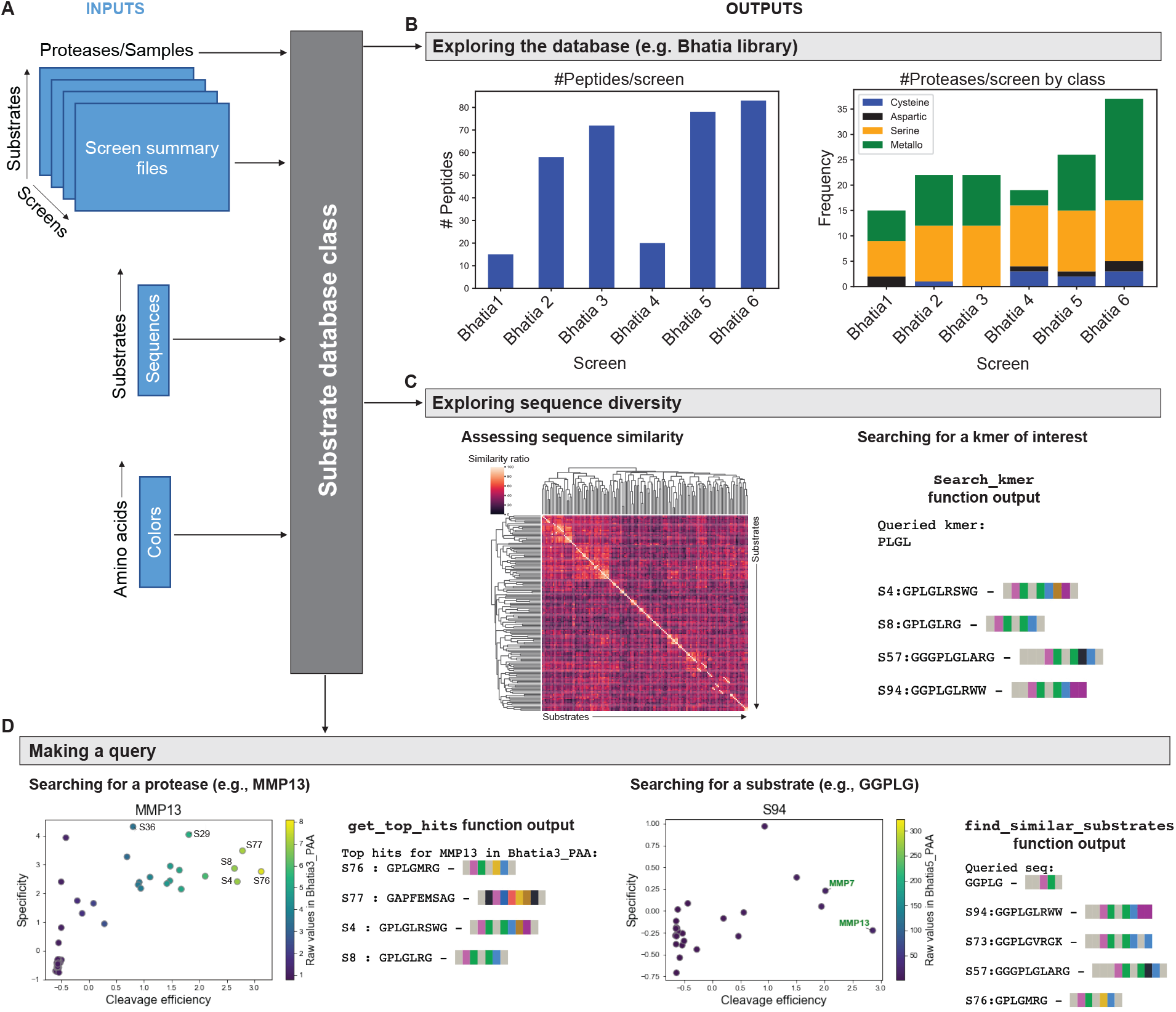
PAA provides infrastructure for queryable databases of enzyme substrates. **(A)** *In vitro* activity screen summary files, a substrate sequence file, and an amino acid color map file provide dataset inputs for a `SubstrateDatabase`. **(B)** Sample use of `SubstrateDatabase` to query a database of 150 unique substrate sequences screened against a diverse set of recombinant proteases. Summary plots of number of substrates and proteases across 6 independent screens comprising the database. **(C)** Metrics of sequence diversity include hierarchical clustering of pairwise sequence similarity scores as well as the ability to search for *k*-mers of interest. **(D)** Sample outputs of querying the database for a protease of interest (e.g. MMP13) and a sequence or cleavage motif of interest (e.g. ‘PLGL’).

To instantiate a `SubstrateDatabase`, the user inputs raw data matrices of activity measurements (i.e., *n* × *k* where *n* is the number of substrates screened, and *k* is the number of conditions, e.g., proteases, assayed) for screens to be included in the database; a file that maps substrate names or labels to their corresponding sequences; as well as an optional file that maps amino acids to different colors based on properties of interest (e.g. hydrophobicity, chemistry, identity) (Fig. 2A). The `SubstrateDatabase` class first identifies overlapping substrates or proteases across multiple screens and aggregates all the data available for each unique substrate and protease into one simple data structure. In this way protease-substrate activity assay data for proteases, substrates, or sequences of interest can be easily and efficiently queried. For example, given a protease of interest as the user query, PAA’s `Substrate-Database` will identify substrates from the database that are most robustly and specifically cleaved. Similarly, given a substrate as the user query, it will identify the proteases that most robustly and/or specifically cleave that substrate. Note that, for the public dataset published as part of PAA, substrate names correspond to names assigned by our group for specific sequences and additionally map to existing nomenclature from previously published work for easy referencing.^12,13,29^

In the absence of an exact match between the query sequence of interest and substrates in the database, `SubstrateDatabase` retrieves the top-*k* substrates most similar to the query sequence, as quantified by different sequence similarity metrics, and returns the *k* sequences as well as the values of these similarity metrics (Fig. 2D). PAA offers two different sequence similarity metrics: the Levenshtein distance similarity ratio and the partial ratio. The former is strictly based on the Levenshtein distance that can be computed as the minimum number of single-character edits (insertions, deletions, or substitutions) required to change one string, or in this case one amino acid sequence, into the other. The partial ratio metric works similarly but instead takes the shortest sequence and compares it with all substrings of the same length. This partial ratio is particularly useful when two substrates contain the same amino acid cleavage motif (e.g., ‘PLG’), but are flanked by different spacers at the N- or C-terminus (e.g., ‘GG’ or ‘GS’ spacers), as they will still be assigned high similarity estimates.

Furthermore, the database also incorporates informative metrics on sequence diversity across substrates (Fig. 2C). Such estimates can be very useful during library optimization to characterize the degree of redundancy and orthogonality between substrates in a given peptide library. Alternative metrics have been recently described by others to achieve similar goals.^30^ To this end, PAA incorporates the function `get_similarity_matrix`, that performs hierarchical clustering of pairwise similarity scores between all substrate sequences in the database and affords a compact visualization of sequence diversity. In addition, the `search_kmer` function allows the user to readily find all substrates in the dataset that contain a given *k*-mer motif of interest, such as the metalloprotease cleavage motif ‘PLGL’. By integrating data from both of these analyses, PAA can help guide library optimization by allowing the user to make inferences about clusters of substrates that may have similar protease cleavage susceptibilities based on similarity scores and known cleavage motifs, as well as by identifying substrates that rank most distinct from others in the library and thus may be favored or disfavored based on the application at hand.

All recombinant proteases included in the database are of human origin. Because overlapping specificities have been identified between protease orthologs,^31^ we anticipate that the sequences provided in this database will prove useful to researchers studying proteases in model organisms. To enable search for orthologous protease genes across species, PAA includes a function, `species-to-species`, that builds off of the comprehensive “Mammalian Degradome Database” ^32^ to facilitate mapping of genes across species of interest (human, chimpanzee, mouse, and rat). For instance, the function will map the human protease ‘*MMP9* ‘ to its mouse ortholog ‘*Mmp9* ‘, while alerting the user that human ‘*CTSL*’ does not have a mouse ortholog.

Furthermore, while here we publish data generated from our group as a public resource and as a prototypical example for the `SubstrateDatabase` data structure, PAA’s functionalities and methods are modular and generalizable to a user’s internally-generated datasets or other datasets of interest. Users can use PAA as a local package for analysis of their own datasets, which will be kept private to the user. The `SubstrateDatabase` interface provides a means by which users can readily query activity screening datasets to nominate candidate substrates and to understand enzyme-substrate cleavage patterns. A guide to the core visualization and analytic functions related to the database can be found at https://github.com/apsoleimany/protease_activity_analysis/tree/master/tutorials. This step-by-step guide showcases how to load and query the protease-substrate database that is published with this work.

### PAA provides modular methods to analyze *in vitro* substrate screens

As highlighted by the degree of information that could be harnessed from our publicly available dataset of protease-substrate activity measurements (Fig. 2), *in vitro* enzyme activity assays are vital to characterizing and understanding protease activities and their dysregulation in disease. ^1,10,33^

Despite the broad prevalence of these assays throughout the protease biology and chemical biology communities, as well as the consideration that such assays will only become larger in size as high throughput screening becomes more common, data analysis remains, to the best of our knowledge, entirely manual. There is a dearth of computational tools to automate the analysis and visualization of enzyme activity datasets. Indeed, all the activity measurements from the 6 recombinant screens presented above were gathered manually, requiring hours of work and incurring higher risk for human error. To bridge this gap, PAA introduces a way to represent, store, and analyze these datasets automatically. The `KineticDataset` data object contains a suite of functions for rapidly preprocessing, visualizing, and analyzing these data (Fig. 3).

**Figure 3:**
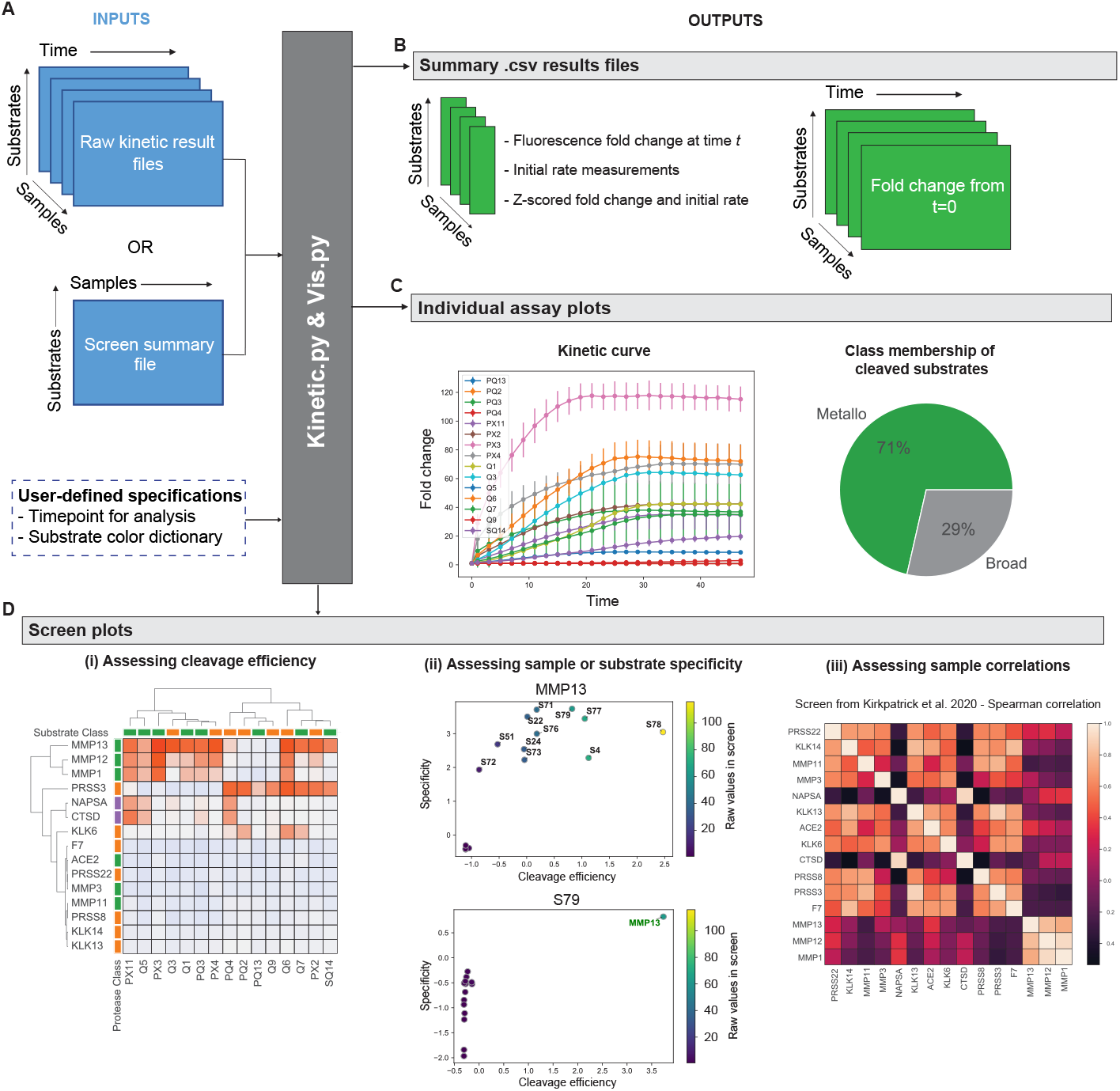
PAA automates analysis of *in vitro* assays of protease activity. **(A)** Fluoresence intensity measurements, together with user-defined specifications such as time points for analysis, are provided as inputs for construction of an instance of the `KineticDataset` class. **(B)** The class automatically generates and saves key output activity measurements, such as initial activity rates and fold increases in substrate turnover as a function of time. **(C)** Retrospective analysis of 15 lung cancer-associated proteases screened against a panel of 14 Forster resonance energy transfer (FRET)-paired substrates,^13^ with representative plots for MMP13 shown, including line plots of fold change intensity over time for each substrate and a pie chart summarizing substrate cleavage susceptibility. **(D)** For the same study,^13^ comprehensive assessment of cleavage efficiency and specificity across recombinant proteases and substrates. (i) Fluorescence fold changes were subject to hierarchical clustering to cluster proteases (vertical) by their substrate specificities and substrates (horizontal) by their protease specificities. (ii) Specificity vs. efficiency (SvE) plots compare standard scores across substrates (efficiency; x-axis) against standard scores across proteases (specificity; y-axis). SvE analysis for the protease MMP13 shows promiscuous activity across substrates (top). SvE analysis for the substrate S79 highlights that it is specifically cleaved by MMP13 relative to other proteases assayed (bottom). (iii) Pairwise correlation analysis of initial rates across all substrates for recombinant proteases in the screen, measured as the Spearman rank correlation coefficient. Heatmap identifies the highest correlation of substrate cleavage between MMP1 and MMP12, amongst all proteases in the analyzed dataset.^13^

The `KineticDataset` class is equipped to take in raw data files from enzyme activity assays (e.g., cleavage of fluorogenic substrates) generated directly by measurement instruments (e.g., fluorimeters; Fig. 3A). Raw files consist of matrices of activity measurements for each sample to be analyzed (i.e., *n* × *t* where *n* is the number of substrates screened and *t* is the number of time-points recorded). The class automatically generates key output activity measurements, including initial rates (*intensity/min*^*−*1^) and fold changes at user-defined timepoints across substrates (Fig. 3B). Functions of `KineticDataset` also compute standard scores (z-scores) across all conditions tested to obtain standardized initial rates and fold changes that are more appropriate for comparing independent assays. Additionally, visualizations can be generated on an instance of `KineticDataset` in order to help interpret the data, such as line plots that depict changes in raw fluorescence intensity and fold change intensity over time for each substrate, and pie charts that summarize substrate cleavage susceptibility to different proteases (Fig. 3C). In this context, the affiliation of a substrate to a particular protease catalytic class is user-defined and can, for instance, refer to the inferred cleavage susceptibility of a substrate by a protease catalytic class (Fig. 3C).

`PAA` also supports inputs from retrospective screens, for which a matrix summarizing cleavage efficiencies across a set of samples may have already been produced (Fig. 3A). Based on these summary matrices, `PAA` can be used to cluster samples (e.g., recombinant enzymes, cell or tissue lysates) of interest based on their activity patterns; to identify substrates that are cleaved with increased specificity by a given sample; and to examine correlations in cleavage patterns across screened samples (Fig. 3D). The specificity vs. efficiency correlation plots (“SvE” plots), in particular, enable identification of optimal protease-substrate pairs from *in vitro* activity assays (Fig. 3D). SvE plots are generated by calculating the z-scores across the screened substrates, which serves as a surrogate metric for cleavage efficiency, and z-scores across the screened proteases, which serves as a surrogate metric for specificity. By plotting these metrics against each other, optimal substrate-protease pairs can be rapidly identified from large sets of screening data by identifying substrates that score high for both of these metrics (Fig. 3D). In addition, annotation by the raw activity measurement values for each protease-substrate pair reflects the absolute cleavage rate of a substrate of interest and overcomes the relative nature of standard scoring. Altogether, these analyses enable rapid assessment of substrate cleavage efficiency and specificity, as well as robust identification of differential or overlapping activity signatures across different enzymes or tissue types (Fig. 3D).

A skeleton framework for the core visualization and analytic functions can be found at https://github.com/apsoleimany/protease_activity_analysis/tree/master/tutorials. The demonstrations and the results presented in this paper feature a retrospective analysis of a previously reported *in vitro* screen of a panel of lung cancer-associated proteases against a panel of 14 peptide substrates^13^ (Fig. 3).

Together, the `KineticDataset` class and the visualization functionality provided by PAA streamline the aggregation, visualization, and analysis of output activity measurements. In particular, these analyses facilitate the assessment of cleavage efficiency and specificity as well as the identification of differential and overlapping activity signatures among different enzymes or tissue types.

### PAA enables machine learning analysis of *in vivo* activity data

Because proteases play critical functional roles in a variety of disease processes, recent years have seen the emergence of new classes of activity-based diagnostics that are engineered to measure the activity of endogenous enzymes at the site of disease and to generate an output signal that can be read out externally.^11,34^ To this end, our group has developed activity-based nanosensors (ABNs), which are probes that detect the activity of aberrant proteases within the body, thereby generating exogenous urinary reporters that reflect the degree of proteolytic cleavage encountered *in vivo*.^12–14,16,35–39^ These nanosensors consist of an inert scaffold whose surface is decorated with peptide substrates, designed to be cleaved by proteases dysregulated in the disease state of interest. Each substrate is marked with a mass-encoded peptide barcode, which is released upon interaction of the nanosensor with target proteases and then concentrated in the urine. Upon collection of urine, the relative concentrations of each reporter are quantified using mass spectrometry. Multiple sensors can be multiplexed in a single *in vivo* administration via barcoding each unique peptide substrate with a different mass-encoded reporter.^5,12–14,16,29^ This multiplexing generates matrices of urinary reporter measurements (i.e., *n* × *k* where *n* is the number of samples and *k* is the number of sensors) on which data analysis can be conducted and diagnostic machine learning classifiers can be trained.

Considering the flexibility afforded by this approach, we implemented a modular framework for the preprocessing and machine learning analysis of these mass-barcoded reporter measurements via the class `SyneosDataset`. Our framework provides a variety of capabilities directly tied to biological, diagnostic, and analytic questions of interest. These capabilities include differential enrichment analysis of reporters between conditions (e.g., identifying which reporters are associated with disease state 1 versus disease state 2); dimensionality reduction via principal component analysis (PCA); binary and multiclass classification; feature, sample, and data specification for all analyses; recursive feature elimination; and associated visualizations.To demonstrate these capabilities, we created a step-by-step guide, published as an implementation tutorial, that recapitulates the findings of previously published work demonstrating the noninvasive detection of localized lung cancer in mice using a 14-plex activity-based nanosensor panel.^13^ For the original study, the authors analyzed the *in vivo* data using unique scripts created specifically for their analysis. In our demonstration, we show how the same raw data can be parsed, analyzed, and visualized using the modular functions available in `PAA` (Fig. 4). We also created additional diagnostic classifiers using the package’s machine learning functions (Fig. 4), thus demonstrating how PAA can be used to derive new insights from existing data.

**Figure 4:**
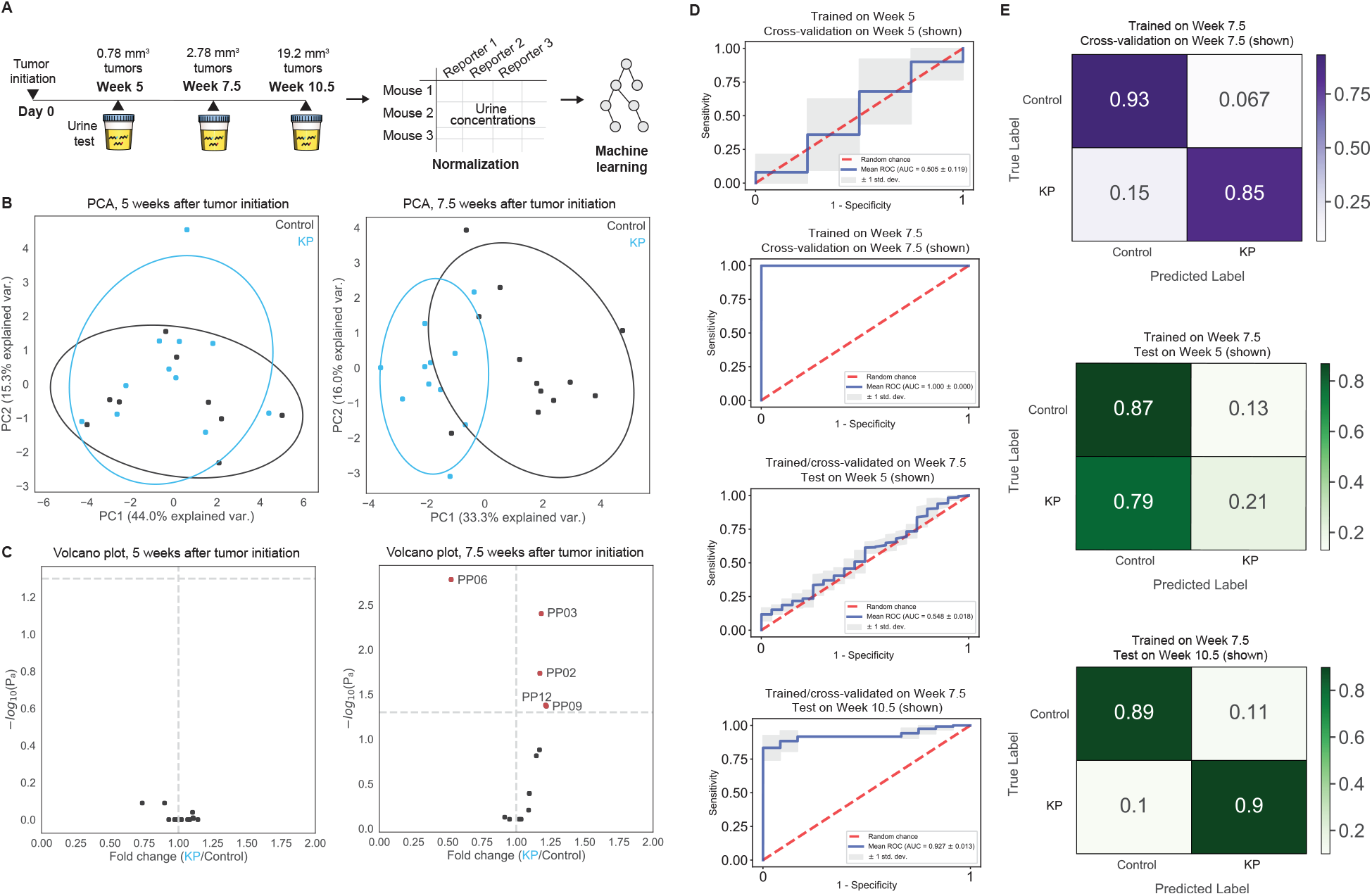
PAA enables automated machine learning analysis of *in vivo* activity data from activity-based nanosensors. **(A)** In a previously published study, activity-based nanosensors were administered at three different timepoints after tumor initiation in a mouse model of lung adenocarcinoma.^13^ Dysregulated protease activity in the cancerous lungs triggered the release of mass-encoded reporters into the urine. The urinary reporter concentrations were measured with mass spectrometry. PAA enables analysis, visualization, and machine learning on these data. **(B**,**C)** PAA automates analysis to visualize differences in reporter enrichment among conditions, such as different sample classes, e.g., wild-type control (Control) and lung cancer (KP) mice, and timepoints, e.g., 5 and 7.5 weeks after tumor initiation in KP mice. **(B)** Principal component analysis (PCA) can reduce the dimensionality of the feature space to discover differential activity signatures across conditions. **(C)** Volcano plots identify nanosensors driving these signatures, by comparing the fold change in reporter concentrations between two classes (x-axis) against their statistical significance (− log_10_(*P*_*adj*_); y-axis). **(D)** PAA evaluates the diagnostic potential of these activity signatures through automated training, validation, and testing of machine learning models, i.e., on the classification of healthy control and KP lung cancer mice. **(E)** Multiclass classifiers can also be trained, tested, and visualized using PAA.

Briefly, in the original study, a 14-plex nanosensor panel was administered into a mouse model of lung adenocarcinoma at 5, 7, and 10.5 weeks after tumor initiation and parallel healthy controls (Fig. 4A). After collecting the urine from each mouse, the urinary reporter concentrations were quantified using mass spectrometry. In the original study, the authors sought to determine the earliest stage at which the nanosensor panel could detect lung cancer. PAA streamlined normalization, statistical analysis, and machine learning of the nanosensor data into a single computational pipeline. We used this pipeline to verify that PAA’s modular workflow could recapitulate the original findings. PAA performed PCA to compare the urinary signals from each disease state across the tested time points (Fig. 4B), and generated volcano plots to visualize which reporters were differentially enriched (Fig. 4C). In the example, one PCA plot (5 weeks after tumor initiation) shows overlap between the two conditions, which is reflected in the volcano plot without any reporters significantly differentially enriched (Fig. 4B). In contrast, the PCA plot showing separation between clusters (7.5 weeks after tumor initiation) corresponds to differentially enriched reporters that drive the separation between groups, as reflected in the corresponding volcano plot (Fig. 4C). All graphs can be generated using one line of code, making such analysis easily accessible and standardized by virtue of using PAA.

The reporter concentrations can then be used to train binary and multiclass machine learning classifiers that can be used to diagnose disease (Fig. 4D-E). PAA is capable of performing classification using support vector machines, random forest algorithms, and linear regression, allowing the user to easily determine the best statistical learning method for their data. The user can also specify which sets of reporters, class labels, or individual labels should be used to train and test the classifiers. In the demonstration, we have shown that the reporter concentrations collected 5 weeks after tumor initiation are unable to learn a representation indicative of lung cancer, whereas by 7.5 weeks, the ABNs are generating an activity signature that can be used to accurately diagnose lung cancer (Fig. 4D-E). More generally, PAA provides a modular, streamlined data analytic workflow for measurements from *in vivo* protease nanosensors, and can readily be applied to new datasets for automated statistical and machine learning analysis.

## Discussion

PAA provides a step towards meeting the need for computational methods to accelerate data analysis in biochemistry, chemical biology, and bioengineering. `PAA` represents a toolkit for users to automate the analysis of protease activity measurements generated *in vitro* through substrate screens or *in vivo* through noninvasive enzyme activity sensors. Here, we focus on analysis of screens against synthetic, fluorescent-quenched peptide substrates (for the former) and of urinary reporter measurements from activity-based nanosensors (for the latter).

However, the modular methods and concepts presented in `PAA` should extend to other datasets, particularly in terms of the tools for analysis of substrate screening data. Additional analytic functions for protease-substrate screening data, such as modeling time to cleavage saturation, prediction of competitive interactions between pairs of peptides,^40^ and identification of optimal substrate sets using principles from information theory,^30^ will expand PAA’s abilities to automate optimal enzyme substrate selection and design. Future work could extend `PAA`’s machine learning capabilities to include neural network models for classification analysis,^41^ methods for assessment of distribution shift and dataset bias,^42–44^ as well as approaches for quantification of predictive confidence.^45–50^

Not only does PAA contain valuable analysis tools, but it also includes a publicly-available database of 150 unique synthetic peptides and their cleavage susceptibilities across a set of 77 distinct recombinant proteases, and provides an interface to query the database for proteases, substrates, or sequences of interest. This resource could be of great interest in the context of protease-cleavable peptide linker nomination to accelerate the development of protease-activated diagnostics and therapeutics. Being implemented and released as a Python package, `PAA` can be further developed and also integrated into larger data analytic workflows in which protease activity measurements play a part. We envision that `PAA` will accelerate analysis workflows for biologists, biochemists, and engineers interested in understanding and leveraging protease activity to better understand, detect, and treat disease.

## Methods

`PAA’s` core relies on `NumPy`, `SciPy`, `Matplotlib`, `pandas`, `seaborn`, and `scikit-learn`. The Python-based implementation allows for flexible use, easy interfacing to machine learning and data analytic packages, and object-oriented programming. `PAA’s` open-source code is maintained on GitHub, is available at https://github.com/apsoleimany/protease_activity_analysis, and published under the MIT license. `PAA` is organized and built as a package for ease of use and to facilitate developer integration.

The demonstrations described in this work are stored as Jupyter notebooks available in the `PAA` repository. These include: (1) analysis and aggregation of *in vitro* screens of recombinant proteases and tissue lysates against synthetic peptide substrates, (2) analysis and machine learning classification of urinary reporter signatures from *in vivo* activity-based nanosensors, (3) querying of protease-substrate database. The datasets used in these demonstrations were generated by our research group and are published together with `PAA`.

All code was written in the Python programming language. The `PAA` package is compatible with Mac OS, Windows, and Linux operating systems.

## Acknowledgement

The authors thank H. Fleming for her thoughtful feedback on this work. A.P.S. acknowledges support from the NIH Molecular Biophysics Training Grant NIH/NIGMS T32 GM008313 and the National Science Foundation Graduate Research Fellowship. C.M.A acknowledges support from “la Caixa” Foundation Postgraduate Fellowship Abroad. M.A. acknowledges support from the National Science Foundation Graduate Research Fellowship. C.S.W acknowledges support from the National Science Foundation Graduate Research Fellowship. S.N.B. is a Howard Hughes Medical Institute Investigator.

